# Scopolamine-induced cholinergic disruption impairs temporal prediction in head-fixed mice

**DOI:** 10.1101/2024.09.06.611607

**Authors:** Yusuke Ujihara, Kota Yamada, Mizuki Yamamoto, Kazuko Hayashi, Koji Toda

**Author notes:** Correspondence: Koji Toda, Ph.D., Department of Psychology, Keio University, 2-15-45 Mita, Minato-ku, Tokyo 108-8345, JAPAN.

## Abstract

Several theories propose that interval timing and memory share common cognitive mechanisms. Pharmacological disruption of the cholinergic system using scopolamine, a muscarinic acetylcholine receptor antagonist, is widely used as a model of cognitive impairment. However, its effects on temporal prediction remain poorly understood. Previous timing studies in freely moving animals are difficult to interpret because motor activity may confound measures of interval timing. Here, we investigated the effects of systemic scopolamine administration on temporal prediction in mice using a head-fixed behavioral paradigm. Mice were trained on a fixed-time schedule task using a peak procedure, in which a 10% sucrose solution was delivered every 10 s through a spout positioned within licking distance. After training, mice developed anticipatory licking responses that peaked around the expected reward time, demonstrating temporal prediction of reward delivery. Systemic administration of scopolamine increased the variability of temporal prediction in a dose-dependent manner without altering the mean peak time. Trial-by-trial analysis further revealed that scopolamine reduced the duration of licking bouts while preserving the total number of licks during peak trials, suggesting impaired maintenance of reward-directed licking behavior rather than altered reward consumption. In a separate open-field assay, scopolamine increased spontaneous locomotor activity and reduced fecal output. These findings demonstrate that cholinergic blockade disrupts the precision of temporal prediction and alters behavioral organization in mice. The increased variability in timing behavior following scopolamine administration may reflect impaired regulation of transitions between behavioral states, providing new insights into the role of cholinergic signaling in temporal cognition.

## Introduction

The ability to predict the timing of environmental events is essential for adaptive behavior, influencing processes such as decision-making, reward anticipation, and learning. Unlike other sensory modalities, temporal information is not encoded by a dedicated sensory receptor or organ but emerges from distributed neural processes. Interval timing, defined as the estimation of durations ranging from seconds to minutes, provides a fundamental mechanism that enables animals to anticipate future events. Although accumulating evidence suggests that cortico-basal ganglia circuits contribute critically to interval timing, the neurochemical mechanisms regulating temporal prediction remain incompletely understood (Buhusi & Meck, 2005; Merchant et al., 2013). Several theoretical frameworks have proposed that interval timing shares common mechanisms with memory processes, particularly in the representation and retrieval of temporal information (Meck et al., 1984; Staddon & Higa, 1993).

The cholinergic system plays a central role in cognitive processes, including attention, learning, and memory. Cholinergic neurons in the basal forebrain provide widespread projections to cortical and subcortical regions and regulate neural plasticity through muscarinic and nicotinic acetylcholine receptors (Ahmed et al., 2019; Deurveilher & Semba, 2011; Hasselmo, 2006; Picciotto et al., 2012). Pharmacological blockade of muscarinic acetylcholine receptors using scopolamine impairs long-term potentiation (Hirotsu et al., 1989; Ito et al., 1988) and disrupts learning and memory performance across species, including rodents (Buresová et al., 1986; Ennaceur & Meliani, 1992; Pitsikas, 2007; Pitsikas et al., 2001; Rudy, 1996, Ozawa et al., 2019; Thonnard et al., 2019), non-human primates (Savage et al., 1996; Oláh et al., 2020; Knakker et al., 2021), and humans (Jones et al. 1991; Wesnes et al., 1991). Importantly, degeneration of basal forebrain cholinergic neurons is a prominent feature of neurodegenerative disorders, including Alzheimer’s disease and dementia with Lewy bodies, and cholinergic dysfunction is strongly implicated in cognitive impairment associated with these conditions (Giacobini, 1990). Consequently, systemic administration of scopolamine has been widely used as a pharmacological model of cognitive dysfunction and cholinergic impairment in experimental studies (Beatty et al., 1986; Collerton, 1986; Ebert & Kirch, 1998; Kopelman & Corn, 1988).

Beyond its well-established effects on memory, accumulating evidence indicates that cholinergic signaling also regulates temporal cognition. Previous studies have reported that systemic scopolamine administration alters timing behavior by reducing temporal precision (Abner et al., 2001; Meck & Church, 1987; Hata & Okaichi, 2001; Balci et al., 2008; Zhang et al., 2019). However, most timing studies have been conducted using freely moving animals, in which motor activity, response initiation, and behavioral strategies may interact with interval timing performance. Although animals can use their own behavioral responses as temporal references (Killeen & Fatterman, 1998; Deverett et al., 2019; Safaie et al., 2020), these movement-related factors complicate the interpretation of whether scopolamine directly affects temporal processing or indirectly alters timing performance through changes in motor output, motivation, or arousal. To overcome these limitations, we recently developed a head-fixed temporal conditioning paradigm that enables precise assessment of temporal prediction while minimizing complex motor sequences associated with freely moving behavior (Inoue et al., 2026; Kaneko et al., 2022; Kaneko et al., 2026; Kawai et al., 2022; Toda et al., 2017; Yamamoto et al., 2022). This approach provides an opportunity to investigate the direct contribution of cholinergic signaling to temporal prediction independently of locomotor confounds.

In the present study, we examined the effects of systemic scopolamine administration on temporal prediction in mice using a head-fixed fixed-time schedule task with a peak procedure. We hypothesized that muscarinic acetylcholine receptor blockade would reduce the precision of temporal prediction rather than simply impair reward-directed behavior or motor performance. To distinguish specific effects on temporal cognition from nonspecific changes in activity, anxiety-related behavior, or autonomic function, we additionally assessed locomotor activity, open-field behavior, and fecal and urine output in freely moving conditions. This experimental approach allowed us to determine how cholinergic disruption influences temporal prediction while dissociating cognitive effects from behavioral and physiological alterations.

## Material and Methods

### Subjects

Eighteen adult male wild-type C57BL/6J mice (age 3.59 ± 2.17 months, range 2.67–6.67 months) were used in this study and maintained on a 12:12 h light/dark cycle. All mice were purchased from Nippon Bio-Supp. Center and bred in the laboratory. Importantly, separate cohorts of mice were used for the head-fixed fixed-time schedule experiment and the open-field experiment. Specifically, six mice (three male and three female; age 2.85 ± 0.03 months, range 2.67–3.00 months) were used exclusively for the head-fixed experiment, whereas twelve male mice were used exclusively for the open-field experiment. All animals were experimentally naive. Mice in the head-fixed experiments had ad libitum access to food, while those in the open-field experiment had unrestricted access to both food and water in their home cages. Body weights were monitored daily, with supplementary water provided post-training as necessary. Experiments were conducted under approximately 75 dB background white noise to mask external sounds, and all the experiments took place during the dark phase of the animals’ light cycle. Following the completion of all experiments, the mice were euthanized humanely using an overdose of isoflurane. All procedures were approved by the Animal Research Committee of Keio University. This study is reported in accordance with the ARRIVE guidelines (https://arriveguidelines.org).

### Surgery

Mice were anesthetized with 1.0–2.5% isoflurane mixed with room air and positioned in a stereotactic frame (942WOAE, David Kopf Instruments, Tujunga, CA, USA) to ensure stable and precise head positioning during head-post implantation. Importantly, no intracranial procedures or brain manipulations were performed. A head post (H.E. Parmer Company, Nashville, TN, USA) was affixed to the surface of the skull along the midline using dental cement (Product #56849, 3M Company, Saint Paul, MN, USA) solely to facilitate subsequent head fixation during behavioral experiments. The skull surface was left intact, and no specific brain region was targeted or assessed by this surgical procedure. Scopolamine was administered systemically via intraperitoneal injection, and the stereotactic apparatus was used exclusively for mechanical stabilization during head-post implantation, not for drug delivery or neural manipulation. Prior to the experiments, mice were group-housed (four per cage), and at least one-week recovery period was allowed following surgery before the onset of the experimental protocols. Post-surgery, each mouse was individually housed in a single cage.

### Behavioral procedure

#### Fixed-Time Schedule Task

Following recovery from surgery (7–14 days), six mice were subjected to water restriction in their home cages (Figure 1A). On the first day of training, the mice were briefly head-fixed and provided with a 10% sucrose solution to habituate them to the experimental environment. Behavioral experiments were conducted in a square chamber (40 cm length × 64 cm width × 60 cm height). Each mouse was positioned on a covered, elevated platform (custom-designed and 3D-printed) with its head fixed using two stabilized clamps holding the sidebars of the headpost (Figure 1B). This head-fixed temporal conditioning paradigm was developed to dissociate interval timing from locomotor behavior (Inoue et al., 2026; Kaneko et al., 2022; Kaneko et al., 2026; Kawai et al., 2022; Toda et al., 2017; Yamada & Toda, 2022; Yamada et al., 2024; Yamamoto et al., 2022). The heights of the tunnel and clamps were adjusted before each session to ensure comfort and stable data recording. A steel drinking spout was positioned in front of the mouse’s mouth, and both the spout and a meshed copper sheet on the stage were connected to a contact touch-sensor (DCTS-10, Sankei Kizai, Tokyo, Japan) that recorded the timing and duration of licking at 1,000 samples/s. Head-fixed mice were allowed to voluntarily lick the spout, with approximately 2 μl of 10% sucrose solution delivered every 10 seconds (Figure 1C). This fixed-time schedule task was adapted from established interval timing procedure used in mice (Inoue et al., 2026; Kaneko et al., 2022; Kaneko et al., 2026; Kawai et al., 2022; Toda et al., 2017; Yamamoto et al., 2022). Sucrose delivery and licking data were recorded using custom Python scripts (version 3.7.7) combined with Arduino UNO and a custom-made relay circuit.

**Figure 1.**
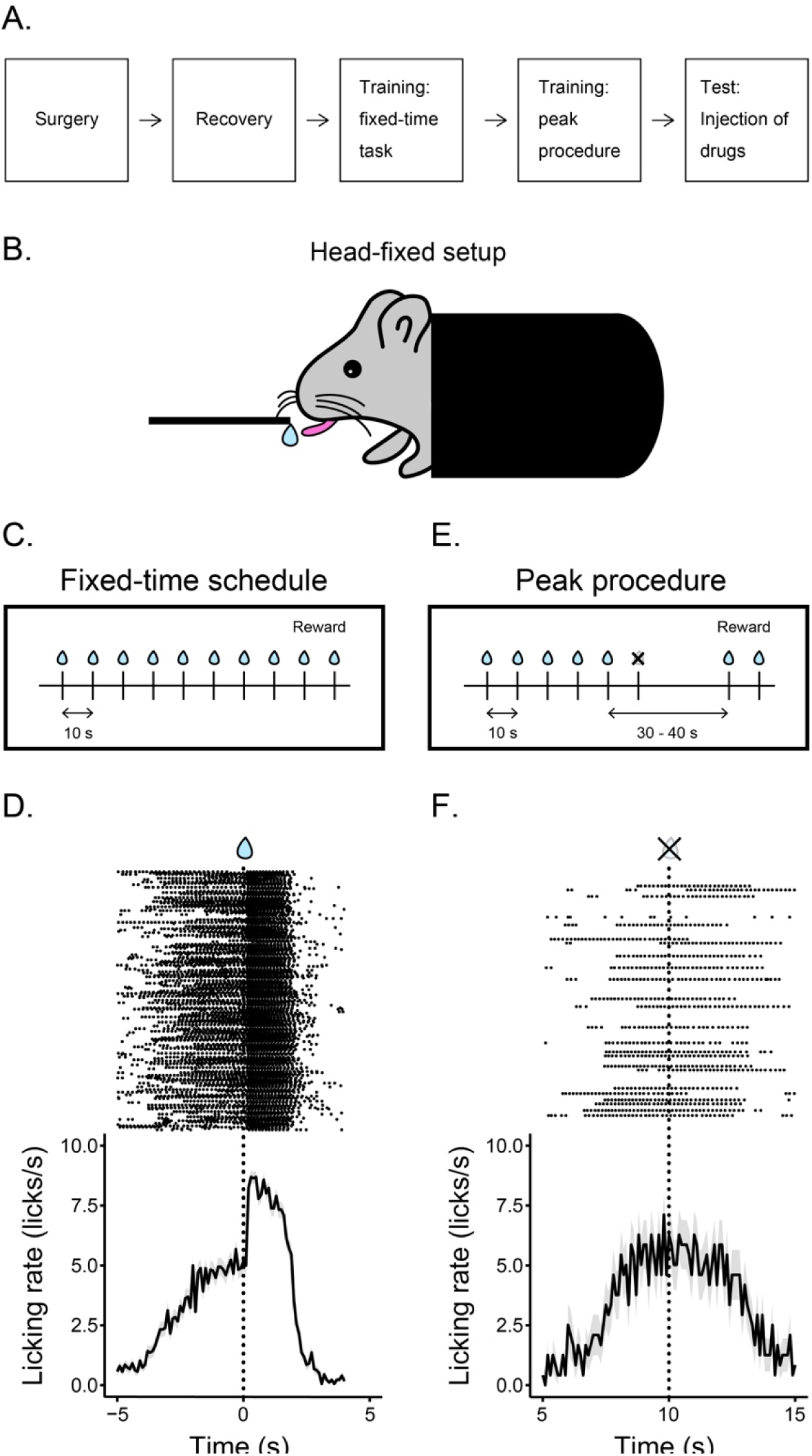
A. Schedule of the head-fixed experiment. B. Head-fixed Behavioral Setup. Mice were head-fixed and positioned in a custom 3D-printed tube. A blunt-tipped needle was situated in front of the mouse for the delivery of a 10% sucrose solution. A custom Python script, in conjunction with an Arduino, controlled and recorded sucrose solution delivery via a solenoid and monitored licking behavior. C. Fixed-Time Schedule. In the fixed-time schedule, a 10% sucrose solution was delivered every 10 seconds throughout the session as a reward. D. Peak Procedure. During the peak procedure, reward delivery was randomly omitted, followed by an extended period without reward delivery. Probe trials lasted 30–40 seconds and were randomly inserted with approximately 20% probability. E. Behavioral Data in Fixed-Time Trials. Raster plots (top) and cumulative Peristimulus Time Histograms (PSTH) (bottom) of licking behavior during fixed-time trials. F. Behavioral Data in Peak Trials. Raster plots (top) and cumulative Peristimulus Time Histograms (PSTH) (bottom) of licking behavior during peak trials.

#### Peak Probe Trials

After the mice demonstrated stable anticipatory licking behavior, probe trials were introduced to assess the accuracy and precision of the mice’s timing of the target interval (Figure 1D). Sessions comprised two types of probe trials: rewarded and non-rewarded. In rewarded probe trials, mice received the same reward as in standard fixed-time trials, namely approximately 2 μl of a 10% sucrose solution delivered at the target interval. Non-rewarded probe trials, or peak trials, allowed the measurement of subjective internal timing in the absence of reward feedback (Balci & Freestone, 2020). During these trials, the interval was extended to three times the target duration (30 seconds), with an additional random duration (mean of 10 seconds, Gaussian distribution). Peak trials constituted approximately 20% of all trials in each session. To maintain task performance, three consecutive rewarded trials were presented following each peak trial. Upon completing training, mice received intraperitoneal saline injections 10 minutes prior to the start of the experiments for 2–3 days to habituate them to the injection procedure. After the saline series, scopolamine was administered systemically via intraperitoneal injection. Mice then received saline injections daily after scopolamine administration until their anticipatory licking behavior returned to normal in the probe trials (2–5 days). Each dose of scopolamine was administered twice on separate days, with saline sessions immediately preceding scopolamine sessions used as controls. The order of injections was randomized.

### Open-Field Test

To assess the effects of scopolamine on spontaneous locomotor activity, an open-field task was conducted using twelve adult male C57BL/6J mice (age 3.96 ± 2.90 months, range 2.76–6.76 months). The procedure for this task was based on our previous work (Inoue et al., 2026; Kaneko et al., 2022; Kaneko et al., 2026; Tamura et al., 2024; Yamamoto et al., 2026). Prior to the experiment, mice underwent habituation to the apparatus and saline injections for 60 minutes per day over three consecutive days. The apparatus consisted of a custom-made white vinyl chloride box (50 cm length × 50 cm width × 50 cm height). Cameras (Logicool HD Webcam C920r, Logicool Co Ltd., Tokyo, Japan) were positioned 106 cm above the floor of the box, and video recordings were captured at 30 frames per second using a Windows PC. During the habituation phase, mice were placed into the open-field box immediately following intraperitoneal saline injections and allowed to explore freely for 60 minutes. During the test phase, mice were placed into the box five minutes after receiving an intraperitoneal injection of either saline or scopolamine. The injection order for saline and scopolamine (0.1, 0.5, and 1.0 mg/kg) was randomized. A minimum interval of two days was maintained between sessions to prevent any carryover effects of scopolamine from influencing subsequent trials. Each open-field test session lasted 60 minutes. White noise (75 dB) was continuously presented throughout the experiment to mask external sounds. After each session, the interior of the box was wiped with 70% alcohol and left to dry for 30 minutes before the next experiment. We employed the open-source visual programming framework Bonsai (Lopes et al., 2015) for computer vision-based tracking of the mice. Video recordings were converted to grayscale, smoothed, and inverted to monochrome. Mice were detected by setting a contrast threshold, and locomotor activity was quantified by tracking changes in the central coordinates of the animals.

### Drugs

To investigate the role of muscarinic acetylcholine receptors in temporal prediction, we administered scopolamine, a muscarinic acetylcholine receptor antagonist. Scopolamine hydrobromide (Nacarai Tesque, Inc., Kyoto, Japan) was dissolved in saline. The doses of scopolamine used in this study (0.1, 0.5, and 1.0 mg/kg) were selected based on prior studies of interval timing in rodents (Balci et al., 2008; Hata & Okaichi, 2001). These studies reported dose-dependent impairments in timing behavior within this range, with lower doses primarily affecting timing precision and higher doses producing more robust cognitive effects. Importantly, these doses have been shown to avoid gross motor impairments, allowing assessment of timing-specific effects.

### Data Analysis

#### Single-session analysis

To calculate peak time and timing precision, a Gaussian function was fit to the normalized peri-stimulus time histogram (PSTH) of licking responses during peak trials for individual sessions. To minimize the potential interference of consummatory licking from previous trials on the peak trial responses, only data within the range of the inter-stimulus interval (ISI) ± ISI/2 (i.e., 5–15 seconds from reward delivery) were used for fitting. The Gaussian function comprises three parameters:

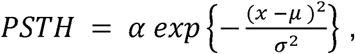

where α represents the peak amplitude, μ denotes the time of peak, σ reflects peak variability and *x* is experimental data. Parameters were estimated using a nonlinear least-squares function with *nls()* function in R (version 3.6.1).

#### Single-trial analysis

To define individual licking bouts during a probe trial, inter-lick intervals (ILIs)—the time elapsed between consecutive licks—were classified into two categories: between-bout ILIs and within-bout ILIs. Bout-and-pause patterns were modeled as a mixture of two exponential distributions (Shull et al., 2001; Killeen et al., 2002):

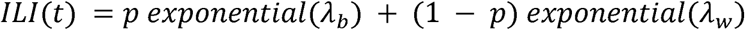

In the equation, *p* denotes the mixing ratio of two distributions, and λ*_b_*and λ*_w_*are parameters for the exponential distributions of between-bout and within-bout ILIs, respectively. The distribution model was fit to the vector of ILIs for each session. Parameters were estimated as the median of MCMC (Markov Chain Monte Carlo) sampling using Stan software (version 2.21.0). We generated 10,000 MCMC samples for the model and data, with 9,000 burn-in periods to discard initial samples, across 4 chains. Convergence was confirmed using the Gelman-Rubin statistic *(R-hat)* with values below 1.1. A No-U-turn sampler was employed to generate MCMC samples. Each ILI was classified into a category based on the model likelihood. For example, if:

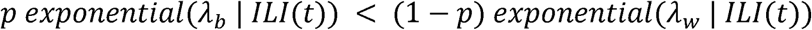

the *ILI* at time t was classified as a within-bout ILI. A cluster of licks adjacent to within-bout ILIs was defined as a bout. Anticipatory licking, consummatory licking, and peak-trial licking were categorized as licks occurring from -4 to 0 s, 0 to 2 s, and 5 to 25 s post-reward delivery, respectively. All analyses and calculations were performed using Stan (version 2.21.0) and R (version 3.6.1).

The position coordinates recorded at 30 frames per second (FPS) were downsampled to 15 FPS by averaging every two consecutive frames. The distance between the position coordinates at time and time −1 was calculated as velocity per frame. Based on this velocity, the mouse’s state was classified into two categories: stopping or moving. A velocity threshold of 0.05 cm per frame was applied, with velocities below the threshold classified as stopping, and those above as moving. The number of bouts initiated during the moving state was calculated by counting the number of transitions from stopping to moving. Bout duration was determined by calculating the total number of moving frames and dividing it by the number of bout initiations.

## Results

### Performance in the Fixed-Time Schedule Task with Peak Procedure

To assess whether mice could predict the timing of reward delivery, they were trained on a 10-second fixed-time schedule task (Figure 1A-C). Head-fixed mice licked a blunt-tipped drinking needle placed near their mouths, which delivered a 10% sucrose solution every 10 seconds. No external cues were provided to indicate the timing of reward delivery. After 5–10 days of training on the fixed-time schedule task, the mice displayed an increasing frequency of licking behavior near the expected reward time, suggesting that they had learned to predict the reward (Figure 1D). Subsequently, peak probe trials were introduced (Figure 1E). In these trials, reward delivery was omitted, followed by an extended period without reward. Probe trials lasted 30–40 seconds and were randomly introduced with a probability of ∼20%. Within a few days, the mice adapted to this peak procedure, with maximal response rates occurring around 10 seconds—the expected reward time (Figure 1F). This indicates that the mice could accurately predict the timing of the reward.

### Effects of Systemic Scopolamine Administration on Interval Timing

To investigate the role of muscarinic acetylcholine receptors in interval timing, we examined the effects of systemic administration of scopolamine, a muscarinic acetylcholine receptor antagonist, on performance in the fixed-time schedule task with peak probe trials in head-fixed mice. Once the mice exhibited the typical response pattern observed in the peak procedure, characterized by a cluster of responses around the anticipated reward time (10 seconds), pharmacological manipulations were initiated. Prior to scopolamine administration, the mice received intraperitoneal saline injections to habituate them to the injection procedure. Systemic administration of scopolamine did not shift the peak response time but significantly increased the variability of temporal predictions in a dose-dependent manner in the averaged data (Figure 2).

**Figure 2.**
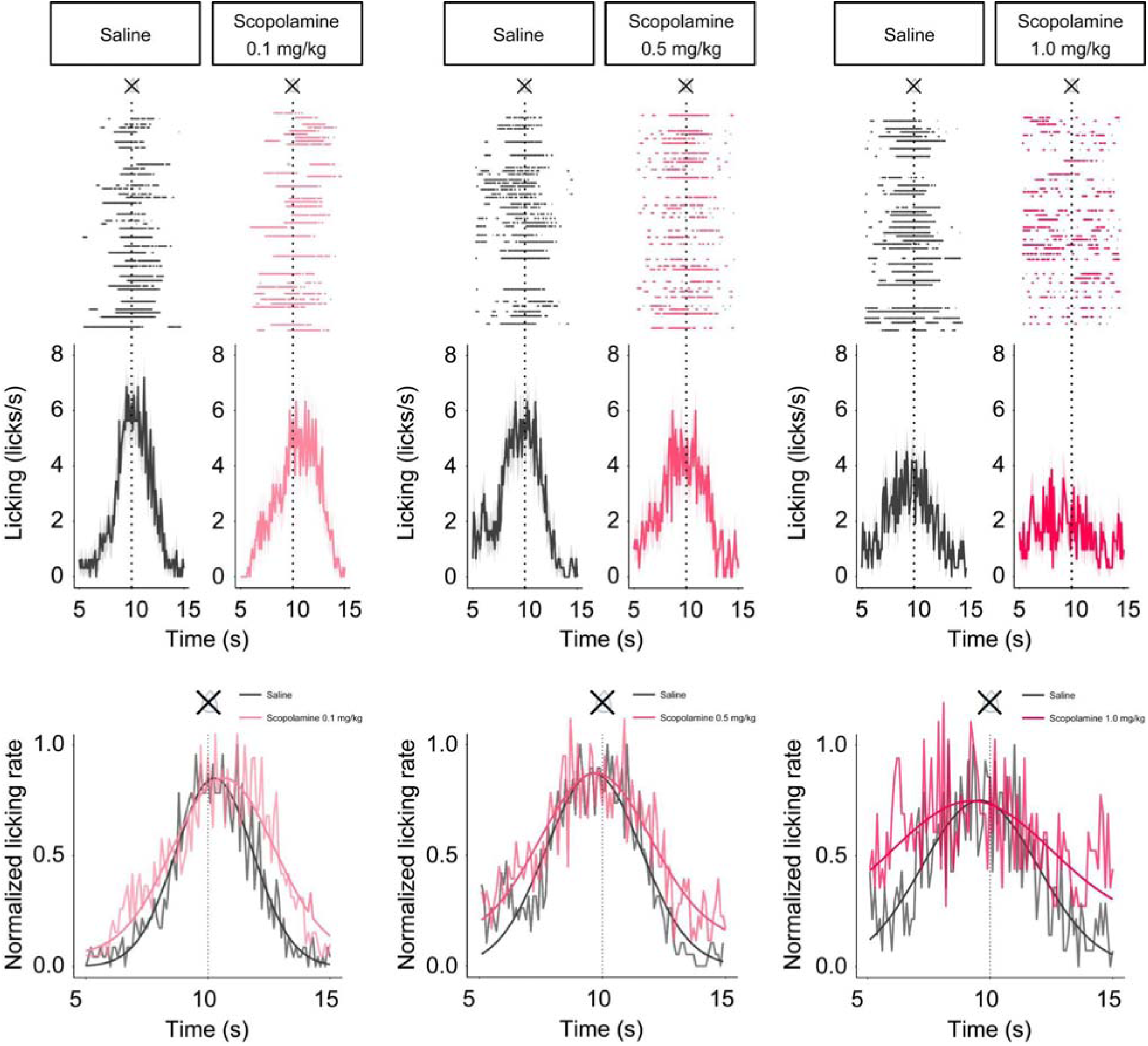
Examples of Raster Data and Peristimulus Time Histograms (PSTH) Following Scopolamine Administration. Raster plots showing licking behavior during peak trials for scopolamine administrations of 0.1 mg/kg (left), 0.5 mg/kg (middle), and 1.0 mg/kg (right) are compared with the saline control (black) sessions (top). Within-bout lickings are represented by colored dots, while between-bout lickings are indicated by gray dots. Corresponding peristimulus time histograms (PSTH) of the data are presented in the middle panels. The bottom panels indicate normalized PSTHs with fitted Gaussian function curves overlaid on the data.

### Quantitative Evaluation of Scopolamine’s Effects on Interval Timing

To quantitatively assess the effects of scopolamine on interval timing, we analyzed peak time and peak variability (Figure 3A). While scopolamine administration at all tested concentrations did not significantly alter the peak time (Figure 3B; F(3, 67) = 0.3267, p = 0.8060, one-way ANOVA), peak variability increased in a dose-dependent manner (Figure 3C; *F* (3, 67) = 12.18, *p* < 0.0001, one-way ANOVA; Saline vs. Scopolamine 0.1 mg/kg: *p* < 0.0001; Scopolamine 0.1 mg/kg vs. Scopolamine 1.0 mg/kg: *p* < 0.0001; Scopolamine 0.5 mg/kg vs. Scopolamine 1.0 mg/kg: *p* = 0.0264, post-hoc Tukey test).

**Figure 3.**
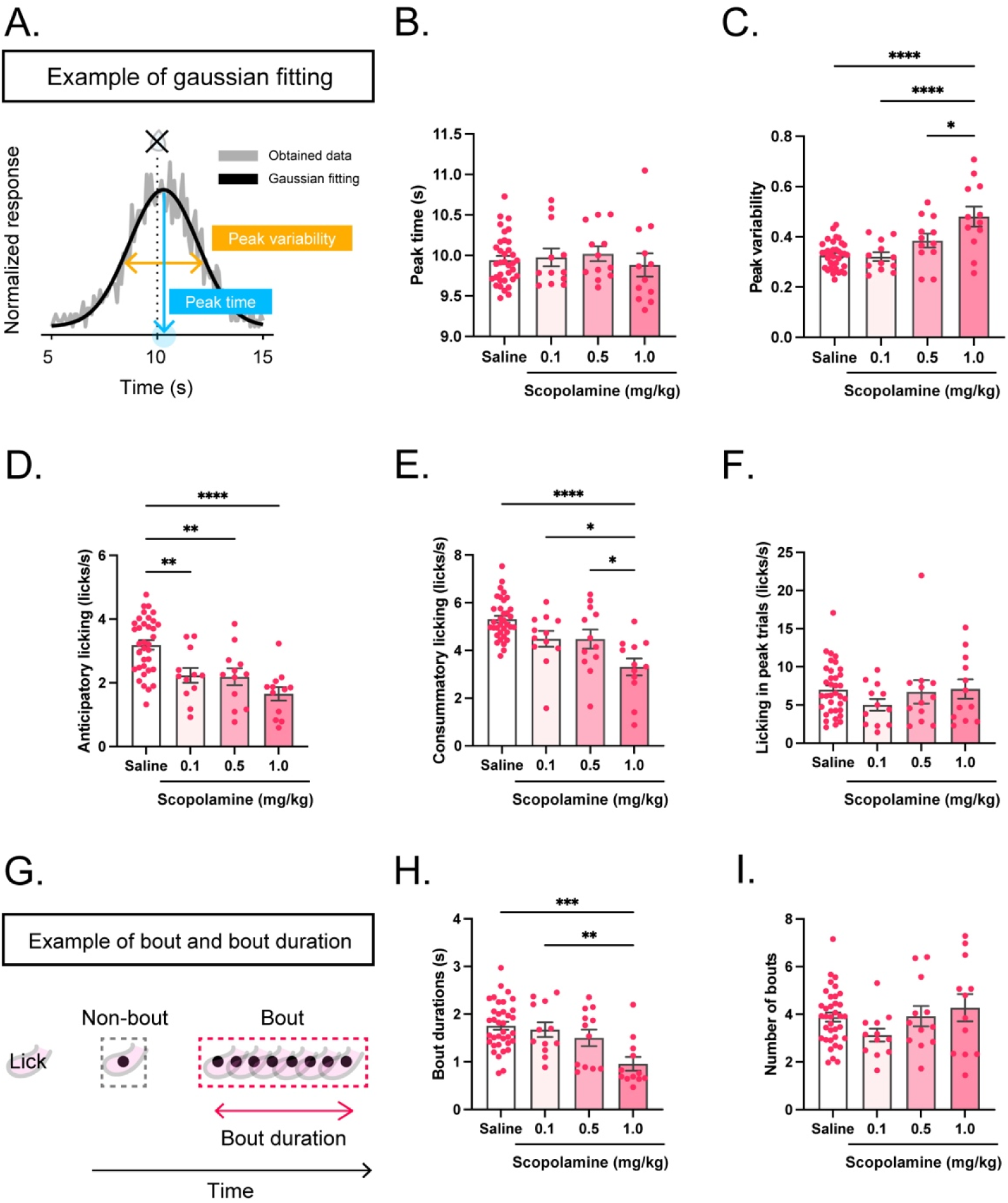
A. Schematic illustration of peak time and variability in peak trials. B. Calculated peak time during peak trials. C. Calculated peak variability during peak trials. D. Total anticipatory licking in peak trials. E. Total consummatory licking in fixed-time trials. F. Total licking in peak trials. G. Schematic illustration of bout and bout length. H. Bout durations during peak trials. I. Number of bouts during peak trials. *Significance levels are indicated as follows: \**p* < 0.05, \*\**p* < 0.01, \*\*\**p* < 0.001, and \*\*\*\**p* < 0.0001. N = 12 (6 subjects).

Next, we examined the impact of scopolamine on licking behavior during interval timing. We classified licking into anticipatory and consummatory phases. Anticipatory licking was defined as licking occurring from -4 to 0 seconds before reward delivery, while consummatory licking was defined as licking from 0 to 2 seconds after reward delivery. Scopolamine administration resulted in a dose-dependent decrease in anticipatory licking (Figure 3D, *F* (3, 67) = 12.19, *p* < 0.0001, one-way ANOVA; Saline vs. Scopolamine 0.1 mg/kg: *p* = 0.0071; Saline vs. Scopolamine 0.5 mg/kg: *p* = 0.0044; Saline vs. Scopolamine 1.0 mg/kg: *p* < 0.0001, post-hoc Tukey test) as well as consummatory licking (Figure 3E, *F* (3, 67) = 10.83, *p* < 0.0001, one-way ANOVA; Saline vs. Scopolamine 1.0 mg/kg: *p* < 0.0001; Scopolamine 0.1 mg/kg vs. Scopolamine 1.0 mg/kg: *p* = 0.0437; Scopolamine 0.5 mg/kg vs. Scopolamine 1.0 mg/kg: *p* = 0.0452, post-hoc Tukey test). However, scopolamine injections did not significantly affect the total number of licks during peak trials (Figure 3F, *F* (3, 67) = 10.83, *p* < 0.0001, one-way ANOVA).

To further explore the effects of scopolamine on interval timing in individual trials, we analyzed the bout structure of licking (Figure 3G). The results showed that scopolamine administration reduced bout duration in a dose-dependent manner (Figure 3H; *F* (3, 68) = 7.179, p = 0.0003, one-way ANOVA; Saline vs. Scopolamine 1.0 mg/kg: *p* = 0.0001; Scopolamine 0.1 mg/kg vs. Scopolamine 1.0 mg/kg: *p* = 0.0070, post-hoc Tukey test), although the number of bouts per trial remained unaffected (Figure 3I; *F* (3, 68) = 1.574, *p* = 0.2037, one-way ANOVA).

### Effects of Intraperitoneal Scopolamine Injection on Locomotor Activity and Excretion

To investigate the effects of the muscarinic acetylcholine receptor antagonist scopolamine on locomotor activity and excretion, we employed an open-field task. Following a 60-minute habituation period in the open-field apparatus over three consecutive days, the experimental phase was initiated. Mice were intraperitoneally injected with saline or scopolamine at doses of 0.1, 0.5, or 1.0 mg/kg, and subsequently placed in the open-field box five minutes after the injection. Scopolamine administration resulted in a significant increase in spontaneous locomotor activity (Figure 4A, left panel; *F* (3, 33) = 8.796, p = 0.0002, repeated measures one-way ANOVA). Increased locomotor activity was observed across all scopolamine doses (0.1, 0.5, and 1.0 mg/kg) compared to the saline condition (Saline vs. Scopolamine 0.1 mg/kg, *p* = 0.0002; Saline vs. Scopolamine 0.5 mg/kg, *p* = 0.0021; Saline vs. Scopolamine 1.0 mg/kg, *p* = 0.0127; post-hoc Tukey test). The effect of scopolamine on locomotor activity was evident shortly after injection and persisted for over 60 minutes (Figure 4A, right panel).

**Figure 4.**
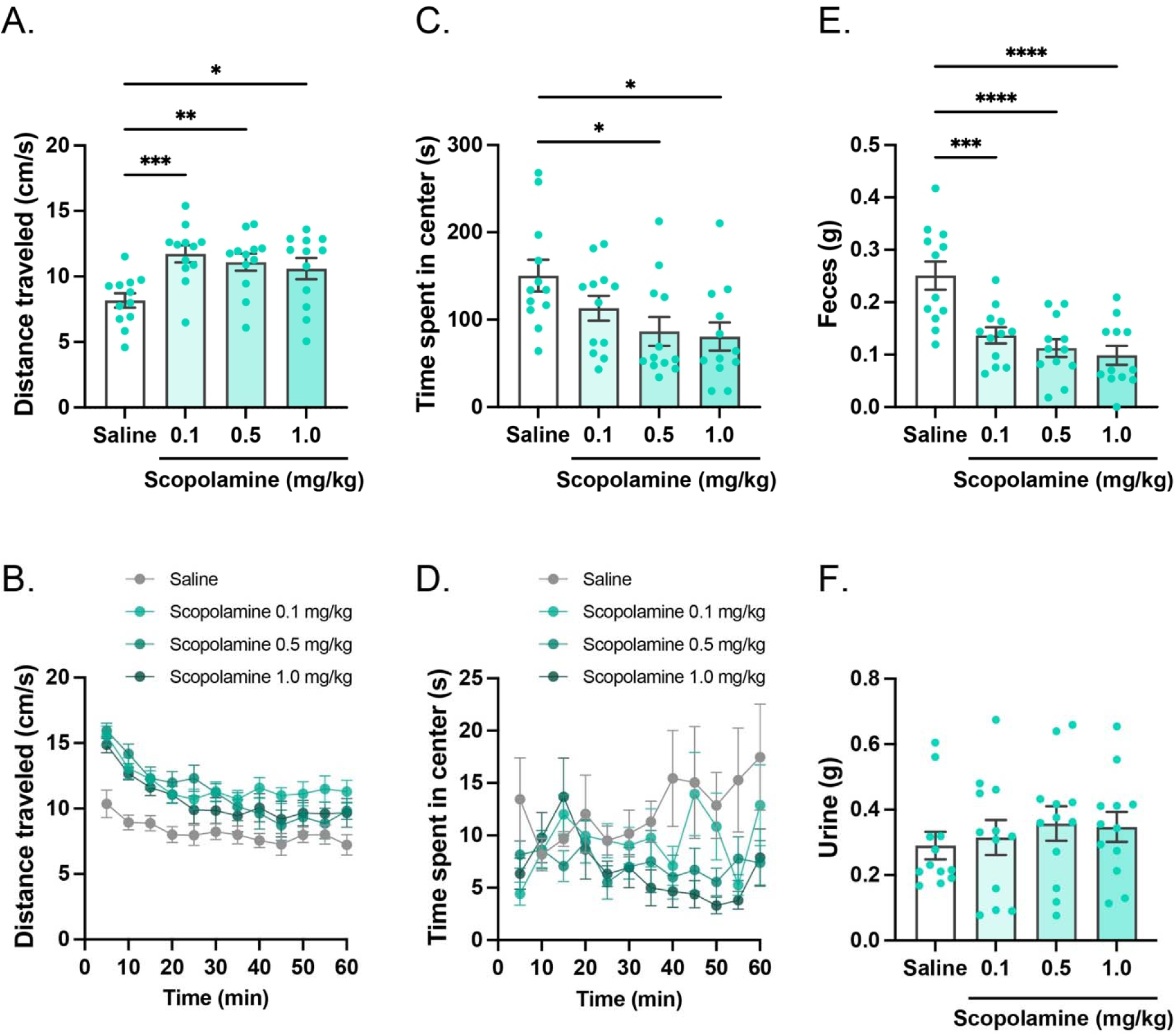
A. Overall locomotor activity from 10 to 60 minutes after the start of the open-field task following saline or scopolamine injection. B. Time course of open-field activity. C. Time spent in the central area of the open-field box from 10 to 60 minutes after the start of the task. D. Time course of time spent in the central area. The central area is defined as the inner 25 cm × 25 cm portion of the box. E. Fecal output collected after 60 minutes of the open-field task. F. Urine output collected after 60 minutes of the task. Error bars represent standard error of the mean (SEM). \**p* < 0.05, \*\**p* < 0.01, \*\*\**p* < 0.001, and \*\*\*\**p* < 0.0001. N = 12.

Additionally, we assessed the time spent in the center of the box as an index of anxiety-like behavior. Scopolamine induced a dose-dependent increase in the time spent in the center area (Figure 4B; *F* (3, 33) = 4.666, *p* = 0.0080, repeated measures one-way ANOVA; Saline vs. Scopolamine 0.5 mg/kg, *p* = 0.0213; Saline vs. Scopolamine 1.0 mg/kg, *p* = 0.0104; post-hoc Tukey test). This effect was observed approximately 40 minutes after the onset of the experiment and continued for over 60 minutes (Figure 4B, right panel).

To evaluate the impact of scopolamine on excretion, a measure of autonomic nervous system function, we quantified fecal and urinary output following the open-field task. Scopolamine administration resulted in a dose-dependent reduction in fecal output (Figure 4C, left panel; *F* (3, 33) = 13.09, *p* < 0.0001, repeated measures one-way ANOVA; Saline vs. Scopolamine 0.1 mg/kg, *p* = 0.0010; Saline vs. Scopolamine 0.5 mg/kg, *p* < 0.0001; Saline vs. Scopolamine 1.0 mg/kg, *p* < 0.0001; post-hoc Tukey test), but did not significantly affect urinary output (Figure 4C, right panel; *F* (3, 33) = 0.5043, *p* = 0.6820, repeated measures one-way ANOVA).

### Analysis of Behavioral Structure in the Open-Field Test Following Scopolamine Administration

To further elucidate the effects of scopolamine on behavioral patterns, we conducted a detailed analysis of spontaneous locomotor behavior during the open-field test (Figure 5). We distinguished between moving and stopping states by calculating movement velocity per frame (Figure 5A). Additionally, we assessed the frequency and duration of movement bouts. Scopolamine administration led to a significant decrease in the number of bouts (*F*(3, 33) = 12.48, p < 0.0001, repeated measures one-way ANOVA; Saline vs. Scopolamine 0.1 mg/kg: p < 0.0001; Saline vs. Scopolamine 0.5 mg/kg: p = 0.0001; Saline vs. Scopolamine 1.0 mg/kg: p = 0.0006; post-hoc Tukey test). Conversely, scopolamine increased the duration of movement bouts (*F*(3, 33) = 9.030, p = 0.0002, repeated measures one-way ANOVA; Saline vs. Scopolamine 0.1 mg/kg: p = 0.0002; Saline vs. Scopolamine 0.5 mg/kg: p = 0.0011; Saline vs. Scopolamine 1.0 mg/kg: *p* = 0.0196; post-hoc Tukey test). These findings suggest that scopolamine decreases the frequency of transitions between moving and stopping states while increasing the duration of movement bouts.

**Figure 5.**
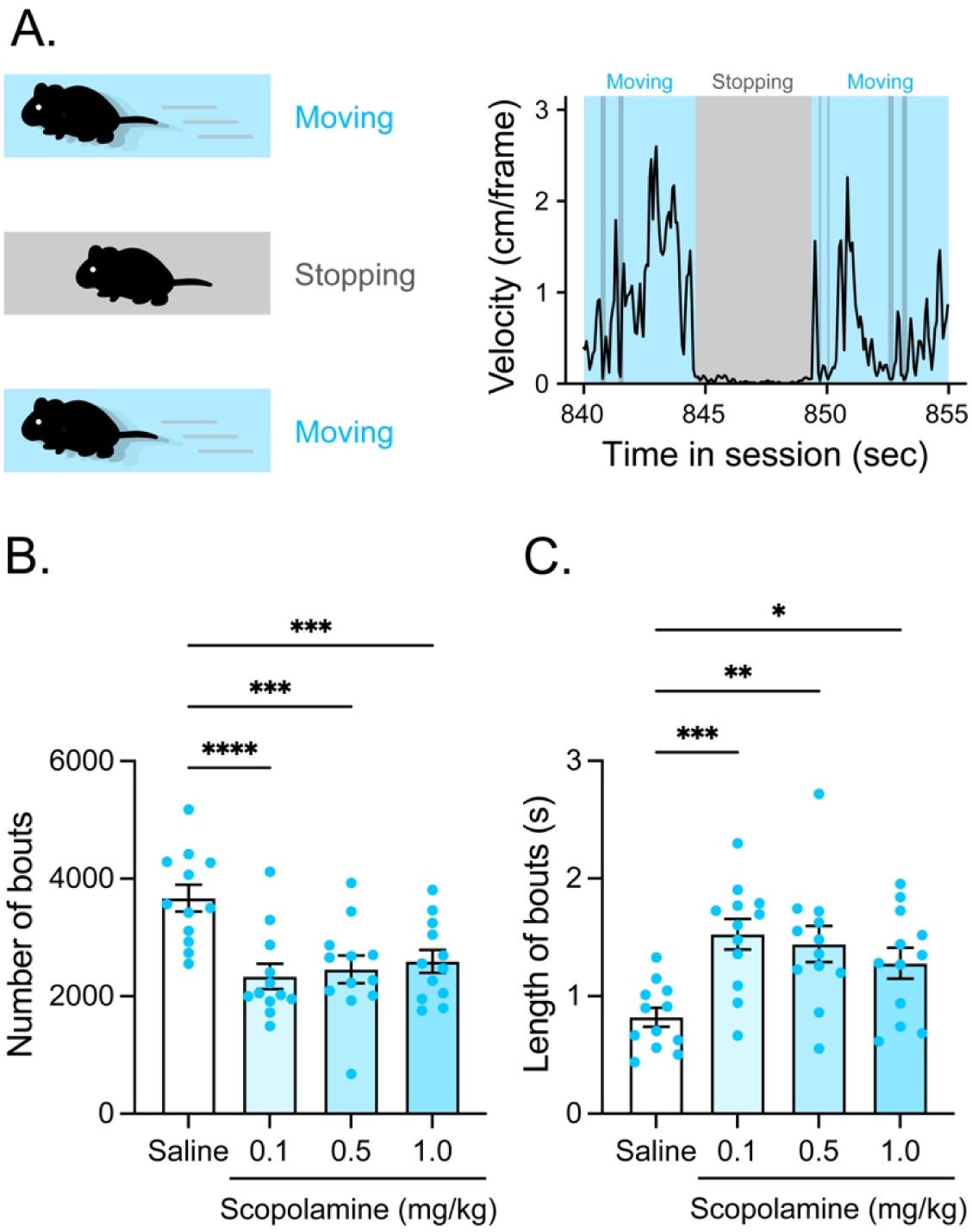
A. We analyzed the structure of spontaneous locomotor behavior in the open-field test. We categorized moving and stopping states by calculating the velocity of movement per frame. In addition, we analyzed the number of the bout and bout durations of the moving state. B. Number of bouts. C. Length of bouts. \**p* < 0.05, \*\**p* < 0.01, \*\*\**p* < 0.001, and \*\*\*\**p* < 0.0001. N = 12.

## Discussion

In this study, we investigated how systemic blockade of muscarinic acetylcholine receptors affects temporal prediction in mice using a head-fixed behavioral paradigm. Mice were trained on a fixed-time schedule task and subsequently tested using a peak procedure, in which they developed anticipatory licking responses that peaked around the expected 10-second reward interval, demonstrating accurate prediction of reward timing. Systemic administration of scopolamine increased the variability of temporal prediction in a dose-dependent manner without altering the mean peak time. Importantly, this effect was observed in a head-fixed paradigm that minimized movement-related confounds associated with freely moving timing tasks. Additional behavioral analyses in a free-moving open-field task revealed that scopolamine increased spontaneous locomotor activity, reduced center exploration, and decreased fecal output. Microstructural analysis further demonstrated that scopolamine reduced moving–stopping transitions and prolonged moving states, suggesting altered regulation of behavioral state dynamics. These findings indicate that muscarinic acetylcholine receptor blockade selectively disrupts the precision of temporal prediction rather than the ability to encode the target interval itself, potentially through impaired stabilization of behavioral states and temporal representations.

Consistent with previous studies using freely moving rodents (Abner et al., 2001; Meck & Church, 1987; Hata & Okaichi, 2001; Balci et al., 2008; Zhang et al., 2019), systemic scopolamine administration increased the variability of temporal prediction. However, interpretation of timing deficits in freely moving animals is complicated by potential contributions from sensory, motor, and behavioral strategy-related factors. Scopolamine is known to induce pupil dilation and impair visual discrimination and attentional processes (Jones & Higgins, 1995; Ennaceur & Meliani, 1992; Parott, 1987). In addition, animals can use their own movement sequences as temporal cues during interval timing tasks, including fixed-interval schedule procedures (Killeen & Fetterman, 1988; Skinner, 1948; Staddon & Simmelhag, 1971; Killeen, 1975). Therefore, the temporal variability observed in previous studies may partly reflect changes in sensory processing, attention, or movement-related strategies rather than a specific impairment of temporal prediction itself. In the present study, the head-fixed fixed-time schedule paradigm minimized these potential confounding factors by eliminating external predictive cues and restricting complex motor sequences during reward anticipation. Under these controlled conditions, systemic scopolamine administration still increased temporal variability without altering the mean predicted interval. These findings indicate that muscarinic acetylcholine receptor signaling contributes to the precision and stability of temporal prediction, rather than simply determining the ability to estimate interval duration.

Our findings extend previous work examining the role of cholinergic signaling in temporal prediction. Using a similar head-fixed paradigm, we previously demonstrated that mice retained the ability to predict reward timing even under conditions in which visual input was disrupted following nicotinic acetylcholine receptor antagonism (Kaneko et al., 2022). In contrast, the present study demonstrates that muscarinic acetylcholine receptor blockade selectively increases the variability of temporal prediction without eliminating the ability to estimate the target interval. These differential effects suggest that muscarinic and nicotinic acetylcholine receptors may contribute distinctively to temporal cognition, with muscarinic signaling playing a critical role in maintaining the precision and stability of temporal representations. Future studies directly comparing muscarinic and nicotinic receptor manipulations using the peak procedure will be important for clarifying the specific contributions of cholinergic receptor subtypes to temporal processing.

A major advantage of the present study is the use of single-trial bout analysis to examine the microstructure of timing behavior. Traditional timing studies often rely on averaged response patterns across multiple trials and sessions. However, such averaging may obscure trial-by-trial behavioral dynamics and generate apparent ramping patterns that do not necessarily reflect the underlying mechanisms (Gallistel et al., 2004a; Gallistel et al., 2004b; Latimer et al., 2015). Consistent with previous studies emphasizing the importance of single-trial analyses in timing behavior (Church et al., 1994; Daniels & Sanabria, 2017; Gallistel et al., 2004b; Toda et al., 2017), we examined the structure of individual licking bouts during the peak procedure. Our analysis revealed that systemic scopolamine administration reduced licking bout duration without affecting the total number of licks during peak trials. These findings indicate that the reduction in temporal precision was not attributable to decreased reward motivation or a general reduction in licking behavior. Instead, scopolamine appeared to impair the ability to sustain continuous reward-directed behavior, suggesting altered stability of behavioral states. The stability of behavioral states may be critical for maintaining internal representations of elapsed time during interval timing. Frequent transitions between behavioral states could interfere with the maintenance of temporal information in working memory, thereby increasing trial-to-trial variability in temporal prediction.

Cholinergic signaling is critically involved in attentional regulation, working memory, and the stabilization of task-relevant neural representations (Klinkenberg & Blokland, 2010). Temporal prediction requires not only the estimation of elapsed time but also the maintenance and updating of internal temporal representations across trials. Therefore, disruption of muscarinic receptor signaling may weaken the neural mechanisms required to sustain temporal information, leading to increased trial-to-trial variability in timing behavior. This interpretation is consistent with the present finding that scopolamine increased variability without altering the average predicted interval, suggesting that cholinergic dysfunction primarily affects the precision and stability of temporal estimation rather than the ability to encode the target interval itself.

Systemic scopolamine administration increased locomotor activity in the open-field task, consistent with previous reports that muscarinic acetylcholine receptor antagonism induces hyperactivity in rodents. (Walters & Block, 1969). Microstructural analysis further revealed reduced transitions between moving and stopping states and prolonged moving periods, suggesting that scopolamine altered the regulation of behavioral state dynamics rather than simply increasing exploratory activity or inducing stereotyped motor patterns. Although we did not directly assess neurotransmitter dynamics, interactions between cholinergic and dopaminergic systems within cortico-striatal and basal ganglia circuits provide a plausible neurobiological framework for these effects. Muscarinic receptor blockade may disrupt cholinergic modulation of dopaminergic signaling, thereby altering behavioral activation and state regulation.

Scopolamine also reduced center exploration and fecal output, indicating effects on behavioral and autonomic states. The reduction in center time may reflect enhanced edge-directed exploration rather than anxiety-like behavior, particularly given previous reports describing anxiolytic and antidepressant-like effects of scopolamine (Drevets et al., 2013; Furey et al., 2010). The decrease in fecal output is consistent with peripheral muscarinic receptor blockade and reduced parasympathetic activity. Although autonomic alterations could potentially influence temporal prediction through interoceptive mechanisms, previous studies suggest that changes in heart rate alone do not necessarily alter temporal prediction (Schwarz et al., 2013). Nevertheless, interoceptive signals may contribute to temporal cognition, particularly under head-fixed conditions in which external cues predicting reward timing are limited. Future studies combining timing tasks with physiological measurements, including heart rate and respiration, will be important for clarifying the relationship between autonomic regulation, interoception, and temporal prediction.

Beyond its peripheral autonomic effects, scopolamine influences central cholinergic systems in brain regions implicated in timing and memory, including the prefrontal cortex, hippocampus, and striatum. These regions constitute key components of the cortico-basal ganglia network that has been strongly implicated in interval timing and temporal prediction (Buhusi & Meck, 2005; Merchant et al., 2013). Cholinergic modulation within these circuits is thought to support cognitive processes underlying temporal prediction by regulating attention, memory, and the stability of task-relevant neural representations. Importantly, temporal prediction in fixed-time and peak procedures is not solely a perceptual process but depends on memory-related mechanisms, including the maintenance and updating of interval representations across trials. Previous theoretical models have emphasized that interval timing relies on working memory and reference memory processes that store and retrieve temporal information (Meck et al., 1984; Staddon & Higa, 1993). In this context, the present finding that scopolamine increased the variability of temporal prediction without altering the mean predicted interval suggests that muscarinic acetylcholine receptor signaling contributes to maintaining the precision and stability of temporal representations rather than generating the basic perception of interval duration itself. Thus, cholinergic dysfunction may impair the consistency with which temporal information is maintained across trials, leading to increased variability in timing behavior.

Several limitations should be considered. First, because scopolamine was administered systemically, we cannot exclude contributions from peripheral muscarinic receptor blockade, including possible effects on autonomic function. Future studies using centrally restricted or peripherally restricted pharmacological manipulations combined with physiological monitoring, such as measurements of heart rate and respiration, will help distinguish central and peripheral mechanisms underlying the effects of scopolamine on temporal prediction. Clarifying the relationship between autonomic fluctuations, interoceptive signals, and temporal precision will be important for understanding the contribution of bodily states to temporal cognition. Second, sex-dependent effects of scopolamine have been reported (Berger-Sweeney et al., 1995; van Hest et al., 1990), but the present study was not designed to systematically evaluate sex differences. Although no apparent sex differences were observed in the head-fixed experiments, the sample size was insufficient for statistical evaluation, and open-field experiments were conducted only in male mice. Future studies should examine sex as a biological variable in cholinergic modulation of temporal cognition. Third, although a within-subject design minimized the influence of inter-individual variability, age was not strictly controlled in some experiments. Since learning and behavioral performance can vary even within the adult age range (Shoji et al., 2016), future studies should carefully control and report animal age to distinguish age-dependent effects from pharmacological effects. Fourth, although our analyses suggest that scopolamine alters behavioral state stability, we were unable to perform detailed classification of behavioral sequences. Future studies using machine learning-based behavioral analysis approaches may provide further insight into how cholinergic signaling regulates behavioral dynamics and temporal prediction. Finally, the present study did not identify the specific neural circuits underlying muscarinic regulation of temporal precision. Future circuit-level investigations using region-specific pharmacological or genetic manipulations will be essential for identifying the brain mechanisms through which cholinergic systems regulate temporal cognition.

## Conclusion

In conclusion, our findings demonstrate that muscarinic acetylcholine receptor signaling plays a critical role in maintaining the precision of temporal prediction by stabilizing internal temporal representations. Systemic scopolamine administration did not alter the average estimate of interval duration but increased trial-to-trial variability, suggesting that cholinergic disruption primarily affects the stability of temporal representations rather than the ability to encode the target interval itself. By combining a head-fixed timing paradigm with behavioral and physiological analyses, this study provides new insight into how cholinergic systems regulate temporal cognition, behavioral state stability, and the neural mechanisms underlying time perception.

## Ethics statement

All procedures were approved by the Animal Research Committee of Keio University (#A2022-129).

## Acknowledgement

This research was supported by JSPS KAKENHI 18KK0070(KT), 19H05316 (KT), 19K03385 (KT), 19H01769 (KT), 20J21568 (KY), 22H01105 (KT), 23H02787 (KT), 23K27478 (KT), 23K22376 (KT), 24H00729 (KT), 24K16869 (KY), and 24KJ0069 (KY), Keio Academic Development Fund (KT), Keio Gijuku Fukuzawa Memorial Fund for the Advancement of Education and Research (KT), Smoking Research Foundation (KT), and HOKUTO Foundation for the Promotion of Biological Science. We thank Kohei Yamamoto and Saya Yatagai for their assistance on animal care.

## Data availability

The data supporting the findings of this study are available from the corresponding author upon reasonable request.

## Code availability

The original codes written for the analyses are available from the corresponding author upon request.

## Contributions

YU and KT designed all the experiments. YU collected all the head-fixed data with the assistance of KY and KT. YU and KT analyzed the head-fixed data with the assistance of KY. YU and KT created the figures for the head-fixed experiment. MY collected all the open-field data. MY and KT analyzed the data of and created the figures for the open-field experiment. YU, KY, MY, KH, and KT discussed the data and commented on the manuscript accordingly. YU and KT wrote the manuscript. KT revised the manuscript.

